# Loss of WNT5 Proteins Reprograms Neutrophils in the Spleen to Provide Protection for DSS-Induced Colitis

**DOI:** 10.1101/2023.01.28.526056

**Authors:** Yi Luan, Jiajia Hu, Qijun Wang, Wenxue Li, Xujun Wang, Rihao Qu, Barani Kumar Rajendran, Hongyue Zhou, Peng Liu, Yu Shi, Yansheng Liu, Jun Lu, Wenwen Tang, Dianqing Wu

**Author notes:** Contributed equally (Yi Luan, Jiajia Hu, Qijun Wang, Wenxue Li, Xujun Wang).

## Abstract

WNT5A and WNT5B are two close homologs, both of which are implicated in the pathogenesis of inflammatory bowel diseases. However, the roles these two proteins play in the disease remain largely uncharacterized. Here, we report that double knockout of *Wnt5a* and *Wnt5b* (*Wnt5* DKO) protects mice from Dextran Sodium Sulfate (DSS)-induced colitis in mice, accompanied with greater splenomegaly, stronger expansion of peripheral myeloid cells, and less colonic CD8^+^ T cell granzyme B expression than those of the control mice. Depletion of neutrophils or splenectomy abrogates the phenotypic differences between *Wnt5* DKO and control mice largely by exacerbating colitis phenotypes and increasing colonic CD8^+^ T cell GZMB expression in the *Wnt5* DKO mice. In addition, neutrophils from the *Wnt5* DKO colitic mice exert stronger suppression of CD8^+^ T cells than those from the control mice in culture. Single-cell RNA sequencing and proteomic analyses indicate that neutrophils from DSS-treated *Wnt5* DKO mice are of hyper-immunosuppressive and hypo-inflammatory characteristics and are distinct from those of DSS-treated control mice as well as myeloid-derived suppressor cells in tumor-bearing mice. Thus, our study reveals that the lack of WNT5 reprograms neutrophils in spleens to limit colonic injury during DSS-induced colitis.

## Introduction

Ulcerative colitis (UC) and Crohn’s disease, collectively known as inflammatory bowel disease (IBD), are highly inflammatory gastrointestinal diseases. IBD affects millions worldwide, particularly in developed countries, and its etiology seems to be very complex and is caused by a combination of environmental factors, defective immune responses, intestinal microbiota, and genetic susceptibility [1–3]. To facilitate the understanding of the disease, mouse models of IBD have been developed. The dextran sulfate sodium (DSS)-induced colitis is the most widely used experimental murine model through the administration of DSS in drinking water. DSS administration leads to erosion of the intestinal epithelium, inflammatory infiltration of the large intestine, and dysbiosis of the intestinal microbiome, which are the features similar to those found in human disease [4, 5]. Although the immune system is a key factor in the pathogenesis of the disease, the involvement of CD8^+^ T cells in the disease seems to be controversial. These contradictions are at least in part due to the different roles of different CD8^+^ T cell subtypes in the disease. Nevertheless, evidence including mouse model studies supports that the conventional cytotoxic CD8^+^ T cells with high granzyme B (GZMB) and interferon-gamma (IFNγ) expression promote both initiation and progression of IBD [6–12].

Wingless-type MMTV integration site family member 5 (WNT5) consists of two members, WNT5A and WNT5B, which share 90% amino acid sequence identity[13]. These WNT5 proteins are involved in diverse physiological and pathological processes, including stem cell self-renewal, cell proliferation, differentiation, migration, adhesion, and polarity[14–16]. *Wnt5a* gene knockout (KO) causes embryonic lethality with significant development defects in mice, whereas *Wnt5b* KO mice show no obvious phenotypes. While evidence indicates WNT5A and WNT5B may have redundant roles in some developmental and physiological processes, including governing the break of left-right symmetry and neuronal specification, evidence also exists for them having distinct functions [15–19]. The WNT5A protein has been linked to intestinal physiology and pathophysiology through multiple regulatory mechanisms. It is involved in crypt regeneration to re-establish homeostasis after intestinal injury [20] and may promote the ability of intestinal dendritic cells (DC) to differentiate naïve CD4^+^ T cells to IFN-γ-producing CD4^+^ Th1 cells in a colitis model [21]. It is not known if WNT5B has any role in intestinal biology and inflammation.

In this study, we investigated the role of WNT5A and WNT5B in the DSS-induced experimental colitis model and found that *Wnt5a* and *Wnt5b* double knockout (*Wnt5* DKO) provided protection to DSS-induced colitis. In addition, the neutrophils from DSS-treated *Wnt5* DKO mice exhibited hyper-immunosuppressive and hypo-inflammatory gene expression characteristics that are distinct from those of myeloid-derived immune suppressors (MDSCs). Furthermore, these cells with increased production of immune suppressive proteins showed enhanced suppression of cytotoxic CD8^+^ T cells and were important for colonic injury protection in the DSS-induced colitis model.

## RESULTS

### *Wnt5* DKO mice were less susceptible to DSS-induced colitis

Homozygous *Wnt5a* KO mice succumb to perinatal lethality due to developmental defects, while homozygous *Wnt5b* KO mice are healthy, develop normally, and are fertile [22]. We generated *Wnt5* DKO mice by crossing *Rosa26Cre^ERT2^* mice, which show ubiquitous expression of Cre/ERT2, with *Wnt5a^flox/flox^* (*Wnt5a^fl/fl^*) and *Wnt5b^-/-^* mice. All these mice had been backcrossed into the C57Bl background prior to the intercrosses. The mice used in this study unless noted otherwise were administrated with tamoxifen at 8 weeks of age, which induces global *Wnt5a* ablation (aka, *Wnt5a^-/-^*) (Fig. S1A). The mice were then treated with 2% DSS delivered in drinking water at 13 weeks of age. The mice lacking both WNT5A and 5B (*Wnt5* DKO) showed less weight loss than the WT control mice after DSS treatment, whereas no significant differences were observed in weight loss after DSS treatment among WT, *Wnt5b^+/-^*, *Wnt5b^-/-^*, and *Wnt5a^-/-^Wnt5a^+/-^* mice (Fig. 1A). Other than weight loss, DSS-induced colitis also displayed pathological symptoms like bloody and pasty stools. These phenotypes were observed and measured as the disease activity index (DAI) to reflect the overall severity of colitis. The severity of colitis was significantly milder in the *Wnt5* DKO mice than that of WT and other *Wnt5* genotype littermates (Fig. 1B). No significant differences in DAI were observed among WT, *Wnt5b^+/-^*, *Wnt5b^-/-^*, and *Wnt5a^-/-^Wnt5b^+/-^* mice (Fig. 1B). Colon length, another colitis-related phenotype, in the *Wnt5* DKO mice was significantly longer than that of WT, *Wnt5b^+/-^*, *Wnt5b^-/-^*, or *Wnt5a^-/-^Wnt5b^+/-^* mice after DSS treatment (Fig. 1C), whereas no difference was observed in colon length before DSS treatment (Fig. S1B,C). Although the loss of WNT5 didn’t alter colon histological structure before DSS treatment, colon tissues in the *Wnt5* DKO mice showed less damaged glands, goblet cells, mucosal architecture, and villa structures than those in the control mice (Fig. 1D-E, Fig. S1D-E). These results together suggest that mice lacking both WNT5A and WNT5B are less susceptible to DSS-induced colitis. Because *Wnt5b* KO alone had no significant impact on DSS-induced colitis or splenomegaly, whereas *Wnt5a^-/-^Wnt5b^+/-^* mice showed trends in these phenotypes, WNT5A and WNT5B are functionally redundant in regulating these phenotypes.

**Figure 1.**
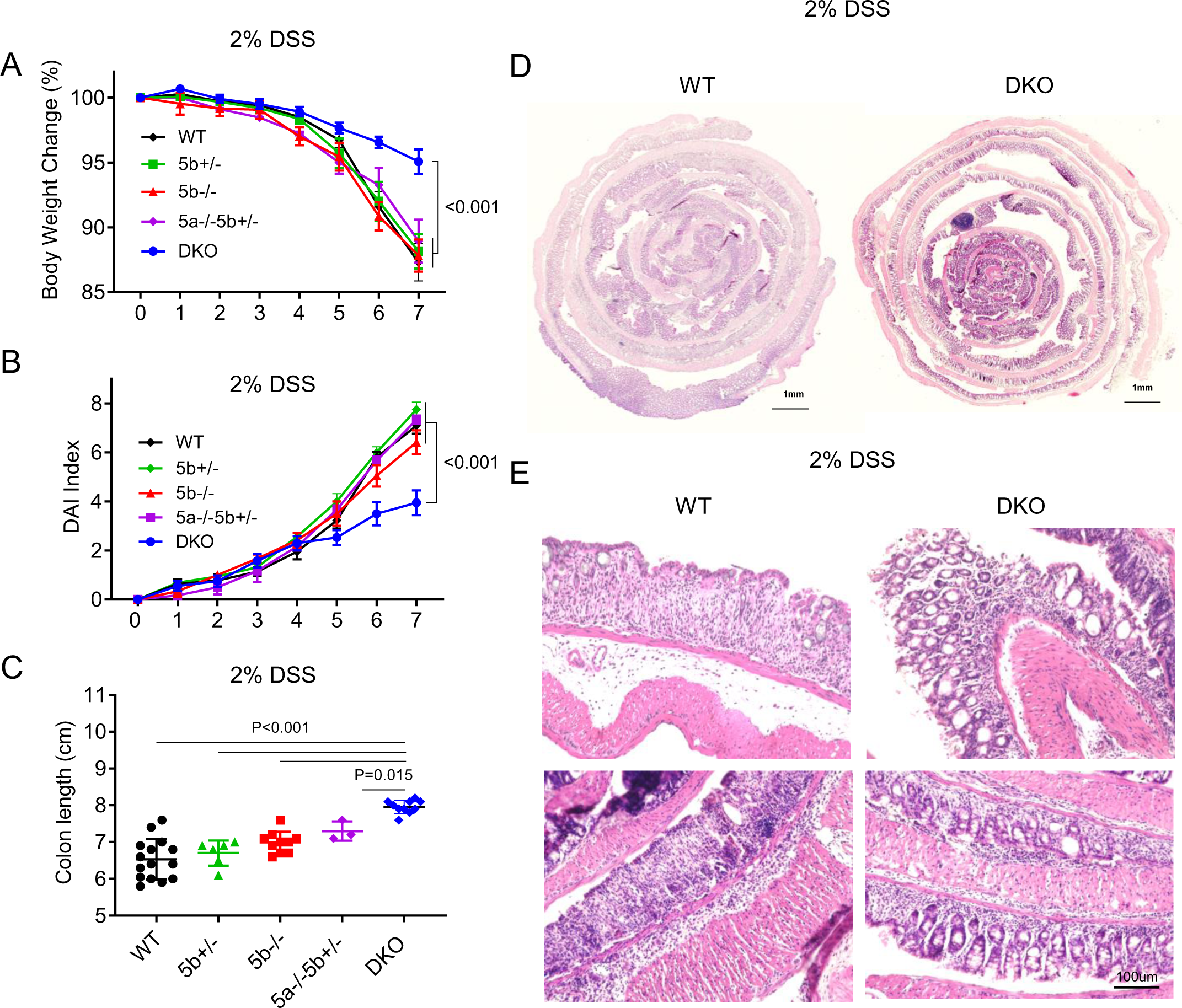
Loss of WNT5 protects mice from DSS-induced colitis. **(A-E)** *Wnt5a* and *Wnt5b* double knock-out mice (DKO, n=9) and various control mice (WT n=13; *Wnt5b^+/-^*, n=7; *Wnt5b^-/-^*, n=8; *Wnt5b^-/-^Wnt5b^+/-^*, n=3) were treated with DSS in drinking water for 7 days. Body weight was monitored daily, and the percentage of body weight loss is shown in **(A)**. Disease activity index was assessed daily and is shown in **(B)**. Colons were collected and analyzed at day 7 of DSS treatment, and colon length is shown in **(C)**. Representative sections of the Swiss rolls from the colons of WT and DKO mice stained with Hematoxylin and eosin (H&E) are shown in **(D)**, and higher magnification views of the colon are shown in **(E)**. In **(C)**, each datum represents one mouse. Results in **(A-C)** are shown as means±sem with P values (two-tailed one-way ANOVA).

### *Wnt5* DKO increases myeloid cell abundance and suppresses cytotoxic T cells in DSS-treated colons

Because immune cells have been shown to play important roles in colitis, we performed flow cytometry analysis of the immune infiltrates in the colons (Fig. S2A-C). Since WT and *Wnt5b^+/-^* mice showed similar colitis phenotypes, we used either WT or *Wnt5b^+/-^* littermates of the DKO mice as the controls in the following experiments. The percentage and the absolute number of neutrophils and monocytes significantly increased in the colon intraepithelial layer (IEL) and lamina propria (LP) of the *Wnt5* DKO mice, while macrophages and dendritic cells remained unchanged (Fig. 2A-D, Fig. S2D-G). CD4^+^ T cells, CD8^+^ T cells, T_reg_ cells, Th_17_ cells, and B lymphocytes were largely unaltered in the colon intraepithelial layer and lamina propria between the *Wnt5* DKO and their WT littermates (Fig. 2E-F & S2H-Q). However, GZMB expression significantly decreased in CD8^+^ T cells in the colons of the *Wnt5* DKO mice (Fig. 2G-H), whereas no significant differences in IFNγ were detected (Fig. S2R). These data together indicate that loss of WNT5 results in an increased presence of myeloid cells and reduced presence of GZMB^+^ CD8^+^ T cells in the colitic colons.

**Figure 2.**
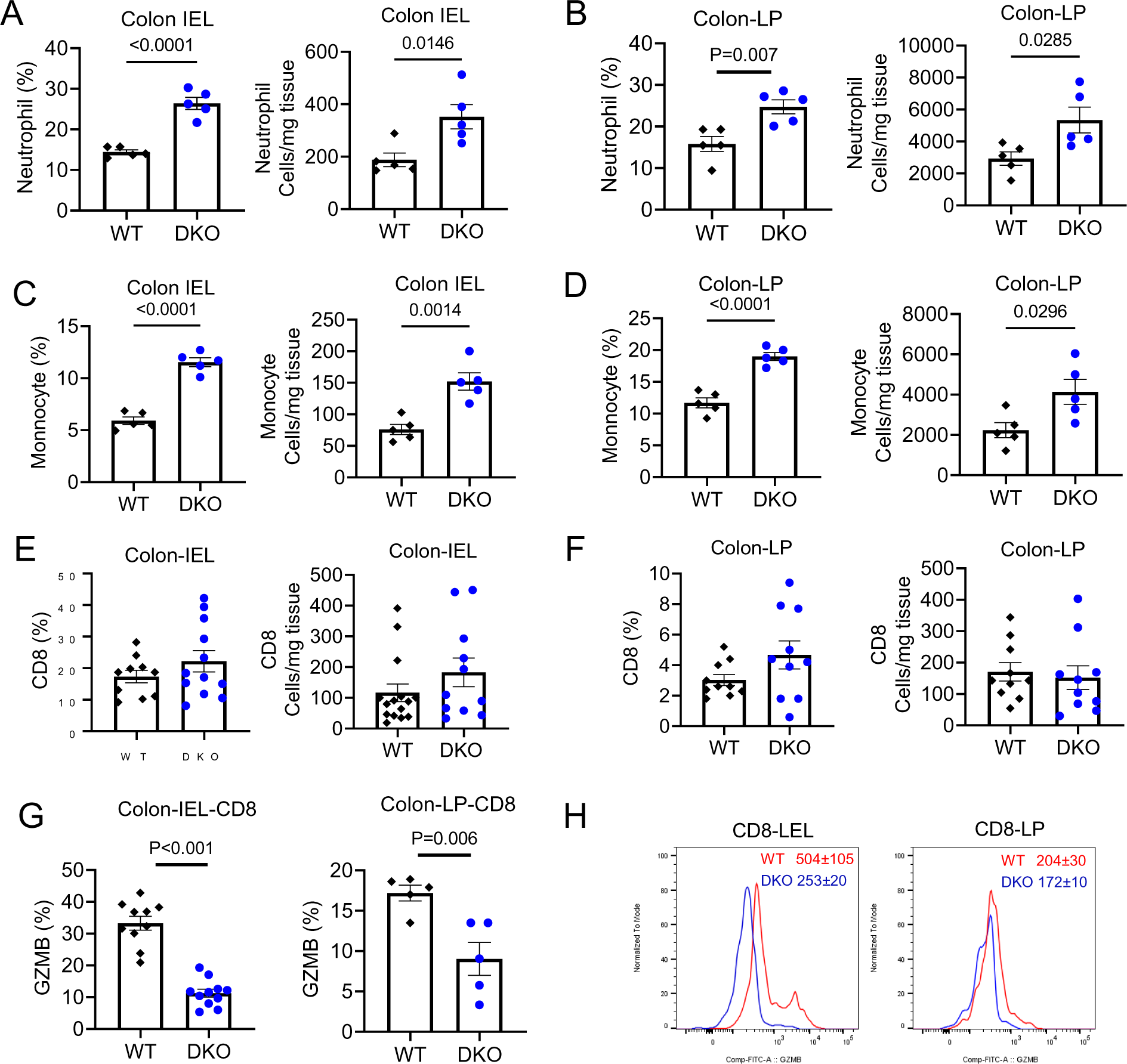
*Wnt5* DKO results in myeloid cell expansion and suppression of GZMB^+^ CD8^+^ T cells in colitic colons. (**A-H**) DKO and WT control mice were treated as in Fig. 1. Cells were isolated from the colon intraepithelial layer (IEL) and laminae propria (LP). (**A-F**) Percentage of leukocyte populations in the CD45^+^ population and their absolute numbers were determined by flow cytometry. (**G**) Percentage of GZMB^+^ in CD8^+^ T cells was also determined by flow cytometry. Mean fluorescence intensity of GZMB in CD8^+^ T cells is shown in (**H**). Each datum point represents one mouse. Data in **(A-G)** are presented as means±sem with P values (two-tailed Student’s *t*-test).

### Colitic *Wnt5* DKO mice develop splenomegaly

During our collection of peripheral lymphoid tissues, we noticed that DSS-treated *Wnt5* DKO mice had significantly enlarged spleens compared to those in the control mice (Fig. 3A & S3A). No difference in spleen weight between the *Wnt5* DKO mice and their control mice was observed without DSS treatment (Fig. S3B). Flow cytometry analysis revealed that the spleens from the *Wnt5* DKO mice had significantly increased percentage and number of neutrophils, monocytes, erythrocytes, and megakaryocytes compared to those of their control mice (Fig. 3B-E & S3C-E). Splenic macrophages showed trends of increase in the *Wnt5* DKO mice, whereas DCs, CD4^+^ T cells, CD8^+^ T cells, and B cells were largely unaltered (Fig. S3F-J).

**Figure 3.**
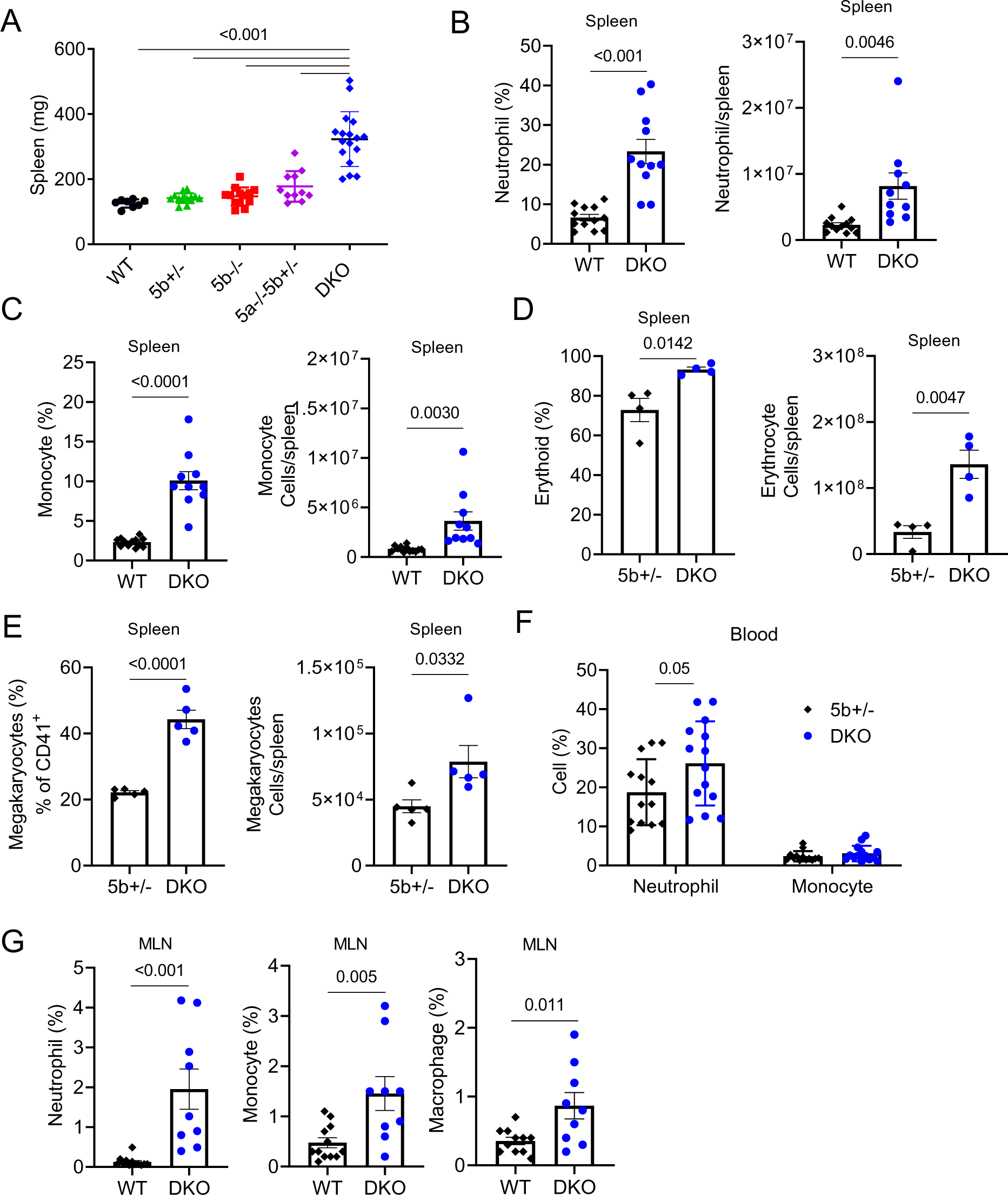
Colitic *Wnt5* DKO mice develop splenomegaly. Mice were treated as in Figure 1, and samples were collected at day7 of DSS treatment. Percentage in the CD45^+^ population and absolute number of splenic, blood, and mesenteric lymph node (MLN) cells were determined by flow cytometry. Data are presented as means±sem with P value (two-tailed one-way ANOVA for **(A)**; two-tailed Student’s *t*-test for **(B)-(G)**). Each datum point represents one mouse.

There were increases in the percentage of neutrophils in the blood and mesenteric lymph node (MLN) of the *Wnt5* DKO mice (Fig. 3F-G), whereas neutrophil percentage remained unchanged in the bone marrow (Fig. S3K). Both monocyte and macrophage percentages increased in the *Wnt5* DKO MLN (Fig. 3G), whereas they did not change in the blood or bone marrow of the *Wnt5* DKO (Fig. 3F, S3K). Additionally, no differences were observed in the percentages of CD4^+^, CD8^+^ T cells, or B cells in the blood, MLN, or bone marrow between the *Wnt5* DKO and control mice (Fig. S3L-N). Taken together, these data indicate that DSS treatment induces greater splenomegaly in the DKO mice accompanied by stronger expansion than the control mice of myeloid cells in the spleen, lymph node, and peripheral blood without obvious effects on other leukocytes.

### WNT5 deficiency enhances splenic extramedullary hematopoiesis upon colitis induction

Splenomegaly is often the result of splenic extramedullary hematopoiesis. Thus, we examined hematopoietic stem and progenitor cells in the spleens of the *Wnt5* DKO and control mice after DSS treatment by flow cytometry. The *Wnt5* DKO mice had significant increases in the percentage and absolute numbers of Lineage^-^cKit^+^Sca1^−^ cells (LK) and Lineage^−^Sca-1^+^c-Kit^+^ cells (LSK) (Fig. 4A & S4A). Although the percentage of long-term hematopoietic stem cells (LT-HSC) and various multipotent progenitor cells (MPP) remained unaltered in the splenic LSK population between the *Wnt5* DKO and control mice, the total number of LT-HSC and MPPs increased in the *Wnt5* DKO spleens (Fig. 4B). In the LK cell compartment, the percentage and the total number of granulocyte and monocyte progenitors (GMP) and megakaryocytes progenitor cells (MKP) increased in the *Wnt5* DKO spleens, which may explain increased neutrophils, monocytes, and megakaryocytes in the DKO spleens (Fig. 4C-D). On the other hand, hematopoiesis showed opposite trends in the bone marrow of the *Wnt5* DKO mice. The percentage and number of LSK cells and the number of LK cells decreased in the DKO bone marrow, while the percentage of LK cells remained unchanged (Fig. S4B). In the LSK cell compartment, the percentage and the total number of LT-HSC and MPP1 showed reductions in the DKO bone marrow. Although the percentage of MPP2 and MPP3 in the LSK population increased in the DKO bone marrow, the total number of MPP2 was unchanged, whereas the number of MPP3 decreased. The number, but percentage, of MPP4 reduced significantly (Fig. S4C). In the LK cell compartment, GMP and MKP were significantly reduced in the *Wnt5* DKO bone marrow (Fig. S4D-E). Collectively, these data indicate that strong splenic EMH accompanied by massive splenomegaly occurred in the *Wnt5* DKO mice after DSS treatment. Meanwhile, bone marrow hematopoiesis subsided in the DKO mice.

**Figure 4.**
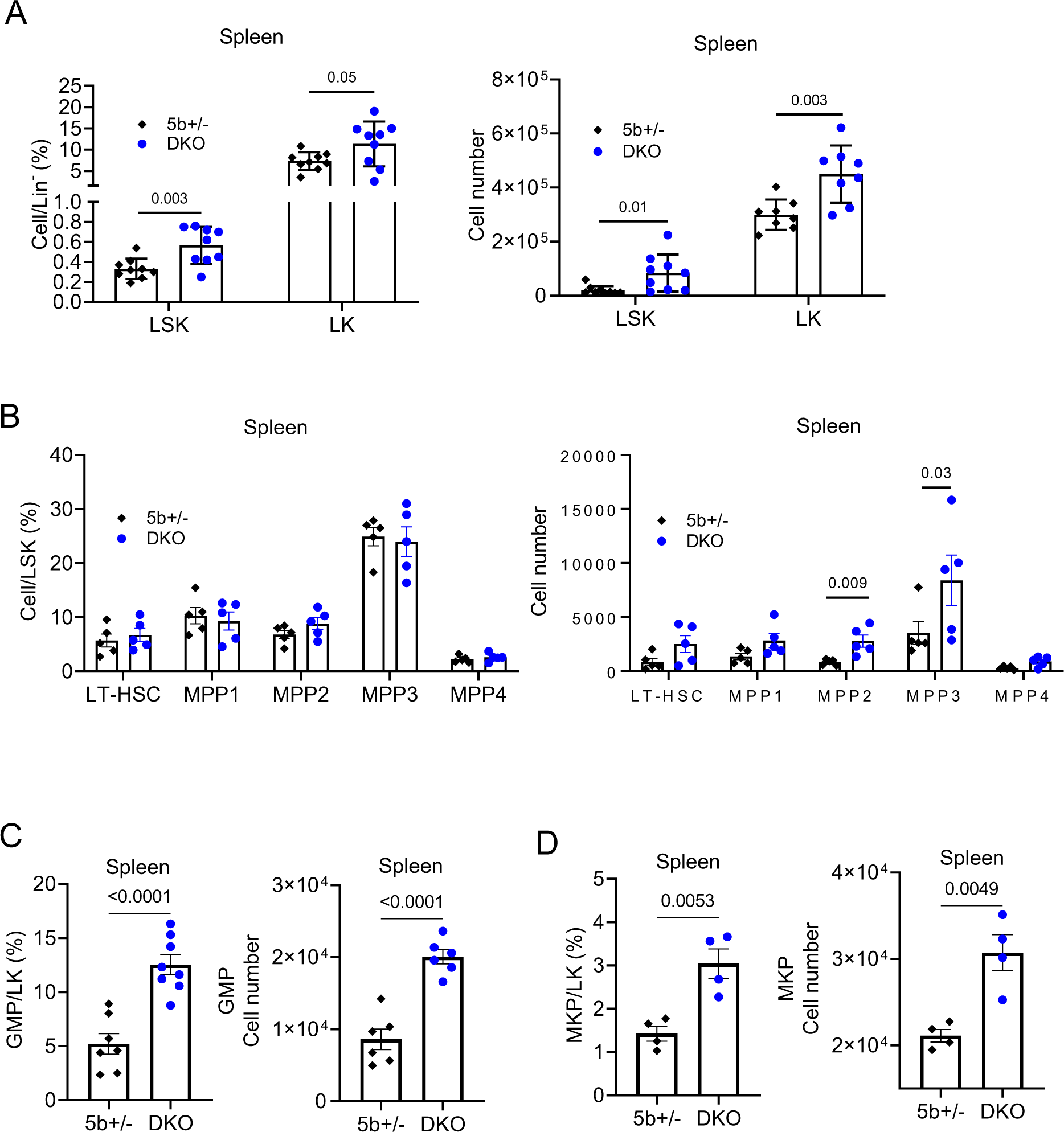
*Wnt5* DKO enhances splenic extramedullary hematopoiesis upon colitis induction. **(A-D)** DKO and WT control mice were treated as Figure 1. Percentage (in Lineage-) and absolute number of LK and LSK **(A)**, percentage (in LSK) and absolute number of LT-HSC, MPP1-4 **(B)**, percentage (in LK) and absolute number of GMP **(C)** and MKP **(D)** in the spleens were determined by flow cytometry. Results in **(A-D)** are shown as means±sem with P values (two-tailed Student’s *t*-test). Each datum point represents one mouse.

### Colitic phenotypes of *Wnt5 DKO* mice depend on splenic neutrophils

The enhanced splenic EMH and expansion of myeloid cells in the spleen, peripheral blood, MLN, and colon in the *Wnt5* DKO mice raised the possibility that the increase in myeloid cells in the DKO mice might be involved in regulating the colitic phenotypes. We examined this possibility by depleting neutrophils by using an anti-Ly6G antibody (Fig. S5A). The administration of the anti-Ly6G antibody effectively depleted neutrophils in the spleen, bone marrow, peripheral blood, and colon (Fig. S5B). Depletion of neutrophils markedly exacerbated colitis phenotypes (body weight, disease index, and colon length) in the *Wnt5* DKO mice and also significantly, but to a lesser extent, aggravate the colitis phenotypes in the control mice (Fig. 5A-C). Importantly, neutrophil depletion abrogated the differences in colitic phenotypes between the *Wnt5* DKO and control mice (Fig. 5A-C). Although splenic neutrophils were successfully depleted, enlarged spleens were still observed in the *Wnt5* DKO mice after anti-Ly6G administration, which might be due to expanded monocytes and erythroid cells in the DKO spleens that were unaffected by neutrophil depletion (Fig. 5D, S5C, D). While the percentages of colonic CD8^+^ T cells as well as CD4^+^ T cells and B cells in the *Wnt5* DKO mice were similar to those of the control mice (Fig. S5E-G), the frequency of colonic GZMB-positive CD8^+^ T cells in colonic CD8^+^ T cells markedly increased upon neutrophil depletion in both control and DKO mice, and again neutrophil depletion abrogated the differences in the frequency of GMZB-positive CD8^+^ T cells between the DKO and control mice (Fig. 5E). These data suggest that neutrophils play an important protective role in DSS-induced colitis and that the neutrophils are responsible for the augmented colitis resistance of *Wnt5* DKO. In addition, these results suggest that neutrophils from the *Wnt5* DKO mice may be more suppressive to CD8^+^ T cells than cells from the control mice. To test this possibility, we performed *in vitro* co-culture of splenic neutrophils with CD8^+^ T cells and found that the neutrophils from colitic *Wnt5* DKO mice showed stronger suppression of granzyme B expression in co-cultured CD8^+^ T cells than those from the WT control mice (Fig. 5F, G). Therefore, neutrophils from *Wnt5* DKO spleens are hyper-immunosuppressive.

**Figure 5.**
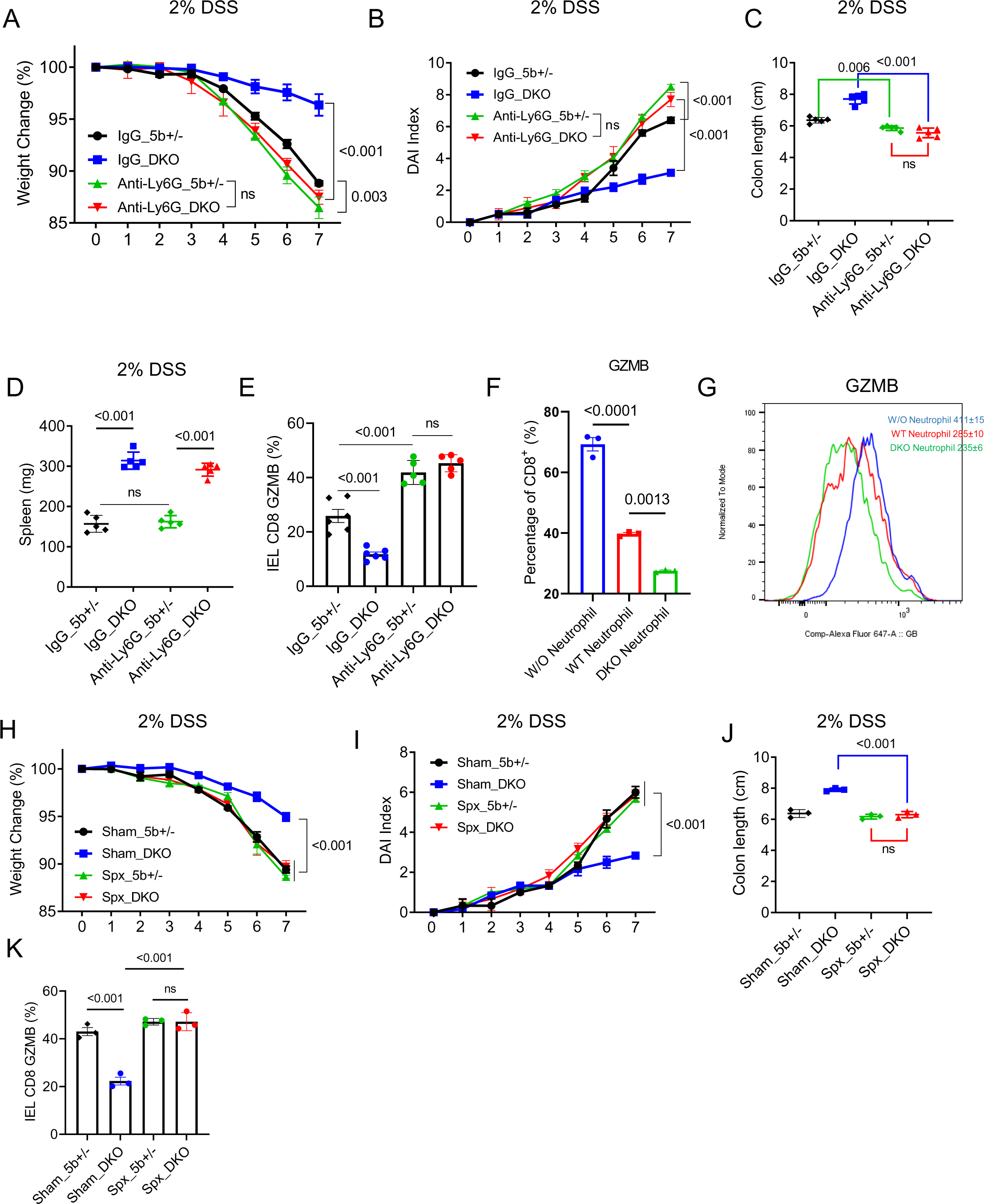
Colitic phenotypes of *Wnt5* DKO depends on neutrophils and spleens. **(A-E)** *Wnt5* DKO (n=5) and the control mice (n=5) were treated with DSS and subjected to neutrophil depletion as shown in Fig. S5A. Colitic phenotypes are shown in (**A-C**), and spleen weight is shown in (**D**). Colon GZMB^+^ CD8^+^ T cells were determined by flow cytometry (**E**). **(F-G)** *Wnt5* DKO and the control mice were treated with DSS, and splenic neutrophils were isolated and co-cultured with isolated CD8^+^ T cells in vitro in the presence of anti-CD3/CD28-beads. Percentage of GZMB in CD8^+^ T cells and representative of flow cytometry traces are shown. **(H-K)** *Wnt5* DKO (n=3) and the control mice (n=3) were subjected to splenectomy before treated with DSS. Colitic phenotypes are shown in (**H-I**), and colon length is shown in (**J**). Colon GZMB^+^ CD8^+^ T cells were determined by flow cytometry (**K**). Data are presented as means±sem with P values (two-tailed two-way ANOVA). Each datum point in **(C)**-**(E)** and **(J)-(K)** represents one mouse. The experiments shown in **(F)** and **(G)** were repeated three times.

The increases in splenic EMH and neutrophil abundance in Wnt5 DKO mice suggest the spleen may be an important source of neutrophils and responsible for the colitis phenotypes of Wnt5 DKO mice. To test this idea, we performed splenectomy in *Wnt5* DKO and control mice right before DSS treatment. The removal of spleens abrogated the differences in the colitis and colonic CD8^+^ T cell phenotypes between *Wnt5* DKO and controls by exacerbating colitis phenotypes and increasing GZMB expression in colonic CD8 T cell of *Wnt5* DKO mice without significantly affecting those of control mice (Fig. 5H-K). These data, together with the data from the neutrophil depletion assay, provide strong support for the conclusion that neutrophils produced by the spleen are largely responsible for the colitis phenotypes of *Wnt5* DKO mice.

#### Neutrophils in colitic *Wnt5* DKO spleen are transcriptomically distinct from MDSCs

To further characterize the effect of *Wnt5* DKO on splenic leukocytes, we carried out single-cell RNA sequencing of splenic CD45-positive cells from DSS-treated WT and *Wnt5* DKO mice, designated as WT-DSS and DKO-DSS, respectively. After quality-control processing, 21745 cells were included in our analysis with an average of 1561 genes per cell profiled, resulting in a total of 21373 mouse genes detected in all cells. Unbiased, graph-based clustering identified 15 clusters and 8 major cell populations (Fig. 6A and S6A, S6B). In line with the flow cytometry data, the percentage of myeloid cells, including neutrophils, monocytic clusters, and their progenitor cells, markedly expanded in the DKO-DSS spleens, while lymphocytes including T cells and B cells remained largely unaltered, compared to those in the WT-Ctrl and WT-DSS spleens (Fig. 6B).

**Figure 6.**
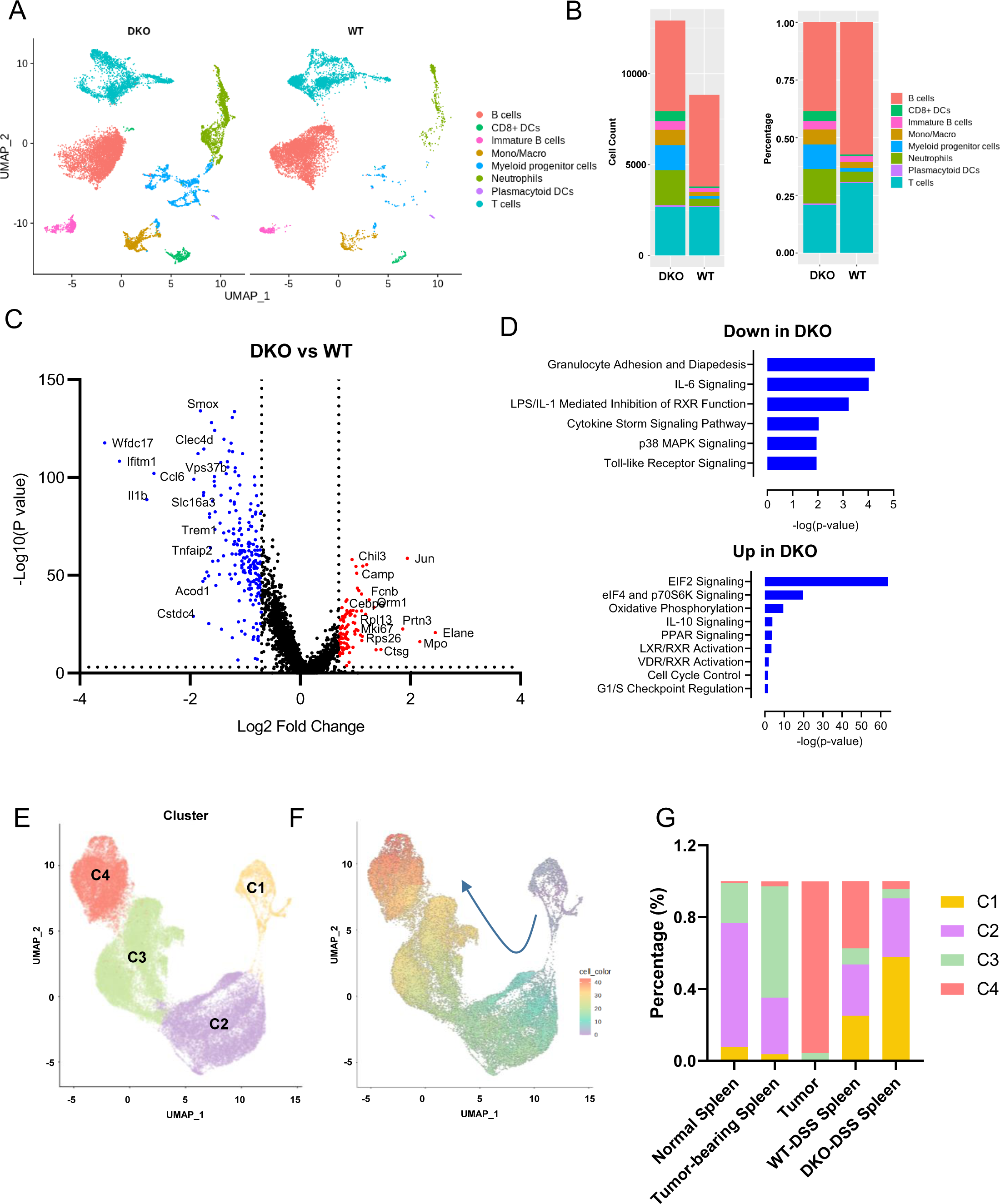
Single cell RNA sequencing analysis of splenic cells. **(A)** UMAP plots of splenic CD45^+^ cells from DSS-treated *Wnt5* DKO (DKO-DSS) and WT (WT-DSS) mice. **(B)** Percentage and total number of each major cell type identified in **(A)**. (**C**) Volcano plot of differentially expressed genes in splenic neutrophils between DSS-treated *Wnt5* DKO (DKO-DSS) and WT (WT-DSS) mice. Significantly upregulated genes (log2 fold change >0.7, P value <0.001) are in red, whereas significantly downregulated ones (log2 fold change < - 0.7, P value <0.001) are in blue. **(D)** Pathway analysis of up or downregulated DEGs between the DKO-DSS and WT-DSS samples. **(E)** UMAP projection of subclusters of neutrophils aggregated from different scRNA-seq analyses. **(F)** Pseudotime plot illustrating the developmental trajectory of the subclusters in (**E)**. **(G)** Percentage of each cluster from **(E)** in different conditions.

Cluster 2, 7, and 14 were identified as neutrophils as these cells were positive for *cxcr2*, *s100a8*, and *ly6g* (Fig. S6C). Differential Gene Expression (DEG) analysis between the WT-DSS and DKO-DSS neutrophils was performed. 194 genes were significantly upregulated, whereas 93 were downregulated in the DKO-DSS splenic neutrophils compared to WT-DSS ones (Fig. 6C, Table. S1). Genes, including *Elane, Mpo, and Mki67*, were upregulated in the DKO-DSS neutrophils. These genes are highly expressed in myeloid progenitor cells[23–25]. In addition, the expression of a number of genes related to ribosomal and protein synthesis, including *Rpl32, Rps18, Rps6, Rps19, Rps26, and Rpl28,* were also upregulated in the DKO-DSS sample. These data indicate that more DKO-DSS splenic neutrophils may be at a proliferative immature stage than the WT-DSS cells. Conversely, genes downregulated in the DKO-DSS splenic neutrophils were related to granulocyte adhesion and proinflammatory responses, among which were *Il1b, Ccl6, Isg15, Cxcr2, and Ccl4.* Ingenuity Pathway Analysis (IPA) based on these DEGs suggested that genes upregulated in the DKO-DSS neutrophils were enriched in the pathways related to EIF2, eIF4 and p70S6K, IL-10, PPAR, and LXR/RXR signaling as well as Cell cycle regulation, whereas the downregulated genes were enriched in granulocyte adhesion, IL-6, p38 MAPK, and Toll-like receptor signaling (Fig. 6D). These data together suggest that splenic neutrophils from the DKO-DSS mice appear to be less mature and hypo-inflammatory than those from the WT-DSS mice.

Given that neutrophils from DSS-treated mice, particularly in the absence of the WNT5 proteins, are suppressive to CD8^+^ T cells, these cells appeared to functionally resemble granulocytic/polymorphonuclear myeloid-derived suppressor cells (PMN-MDSCs). PMN-MDSCs were characterized in tumor-bearing mice and human patients and are primarily produced in spleens via EMH [26–28]. These cells were further characterized recently by single-cell RNA sequencing [26, 27, 29]. We thus wanted to compare the transcriptional characteristics of immune suppressive neutrophils induced by colitis with those of PMN-MDSCs in tumor-bearing mice. We aggregated our single-cell RNA sequencing data with those described in [26, 27, 29] and extracted cells based on PMN cell markers *Ly6g, Cxcr2,* and *S100a8* (Fig. S6D). Unsupervised clustering further partitioned these neutrophils into four clusters, C1-C4 (Fig. 6E and S6E). Genes, including neutrophil progenitor marker (*Elane),* cell cycle and proliferation markers *(Mki67* and *Cdk8)* and various ribosome-related genes related to translation, were enriched in Cluster C1, suggesting that cells in the C1 cluster consist mainly of neutrophil progenitors with proliferative characteristics. The C2 and C3 cluster appeared to represent more mature neutrophils than the C1 cluster as genes related to leukocyte migration, and activation, including *Camp, Mmp8, Itgb2, Csf3r,* and *Cxcr2,* were enriched in these clusters. The C4 cluster seemed to be the most mature population with high expression of proinflammatory cytokine and chemokine genes---*Il1b, Ccl3, Ccl4,* and *Cxcl2*. In addition, many genes, including *Ler3, Nfkbia, Isg15,* and *Il1r2,* related to inflammatory response, cytokine production, and programmed cell death were highly enriched in the C4 cluster (Fig. S6E-F, Table S2, S3). Unsupervised pseudotemporal analysis supported the conclusion that Cluster 1 represented the earlier stage of neutrophil development, which progresses to Clusters 2 and 3, and then Cluster 4 (Fig. 6F). Next, we segregated and compared cells of normal spleens, tumor-bearing spleens, tumors, WT-DSS spleens, and DKO-DSS spleens (Fig. S6G) and found that neutrophils from normal and tumor-bearing spleens mainly belonged to clusters C2 and C3, whereas neutrophils from the tumor tissues were almost exclusively enriched in the C4 cluster (Fig. 6G, S6G). Neutrophils from the WT-DSS spleens were more evenly distributed in clusters C1, C2 and C4, suggesting that DSS-induce colitis triggered a different distribution pattern of neutrophils in the spleen (Fig. 6G, S6G). Strikingly, neutrophils from the DKO-DSS spleens showed a very distinct distribution pattern from those mentioned above, and the majority of the cells were enriched in the C1 cluster with a few in the C2 cluster (Fig. 6G, S6G). As the C1 cluster represents less mature neutrophils, it was consistent with the earlier conclusion that development/maturation of splenic neutrophils seems to be retarded in the absence of Wnt5. Of note, the majority of cells in the C1 cluster do not express neutrophil progenitor marker gene *Elane* (Fig. S6H), and the pseudotemporal analysis with *Elane* expression as the start point of neutrophil differentiation indicates that most of the C1 cluster cells, which the majority of *Wnt5* DKO cells belong to, do not lie in between *Elane*-positive pronator cells and more mature neutrophils in Cluster 2 (Fig. S6I). Thus, *Wnt5* DKO may reprogram the developmental process of neutrophil progenitors.

### Upregulation of immunosuppressive proteins in circulating DKO-DSS neutrophils

To better understand how neutrophils from the DKO-DSS mice are more immunosuppressive, we performed proteomic profiling of circulating neutrophils of the DKO-DSS and WT-DSS mice. The circulating neutrophils are the closest to the effector neutrophils infiltrated in the colon and can be obtained at a sufficient quantity for proteomic analysis. The proteomic analysis revealed 21 proteins significantly upregulated and 400 downregulated in the DKO neutrophils compared with control neutrophils (Fig. 7A, Table S4). Pathway analysis indicated that anti-inflammatory related pathways, including the PPAR and PPARa/RXRa activation pathway, were upregulated in the DKO neutrophils, whereas pathways related to neutrophil pro-inflammatory activation, including cholesterol biosynthesis, oxidative phosphorylation, neutrophil extracellular trap signaling, and IL-6 signaling, were significantly downregulated in the DKO neutrophils (Fig. 7B). These data, in line with the gene expression profile of splenic neutrophils, suggest that DKO neutrophils are less pro-inflammatory. The proteomic analysis also showed that immune regulatory proteins, including CD101, CD47, and Thbs1, were upregulated in the DKO neutrophils. These proteins have been shown to be involved in T-cell suppression[30–34]. The flow cytometry analysis validated the upregulation of these proteins in the circulating neutrophils from the DKO-DSS mice compared to those from the WT-DSS mice (Fig. 7C).

**Figure 7.**
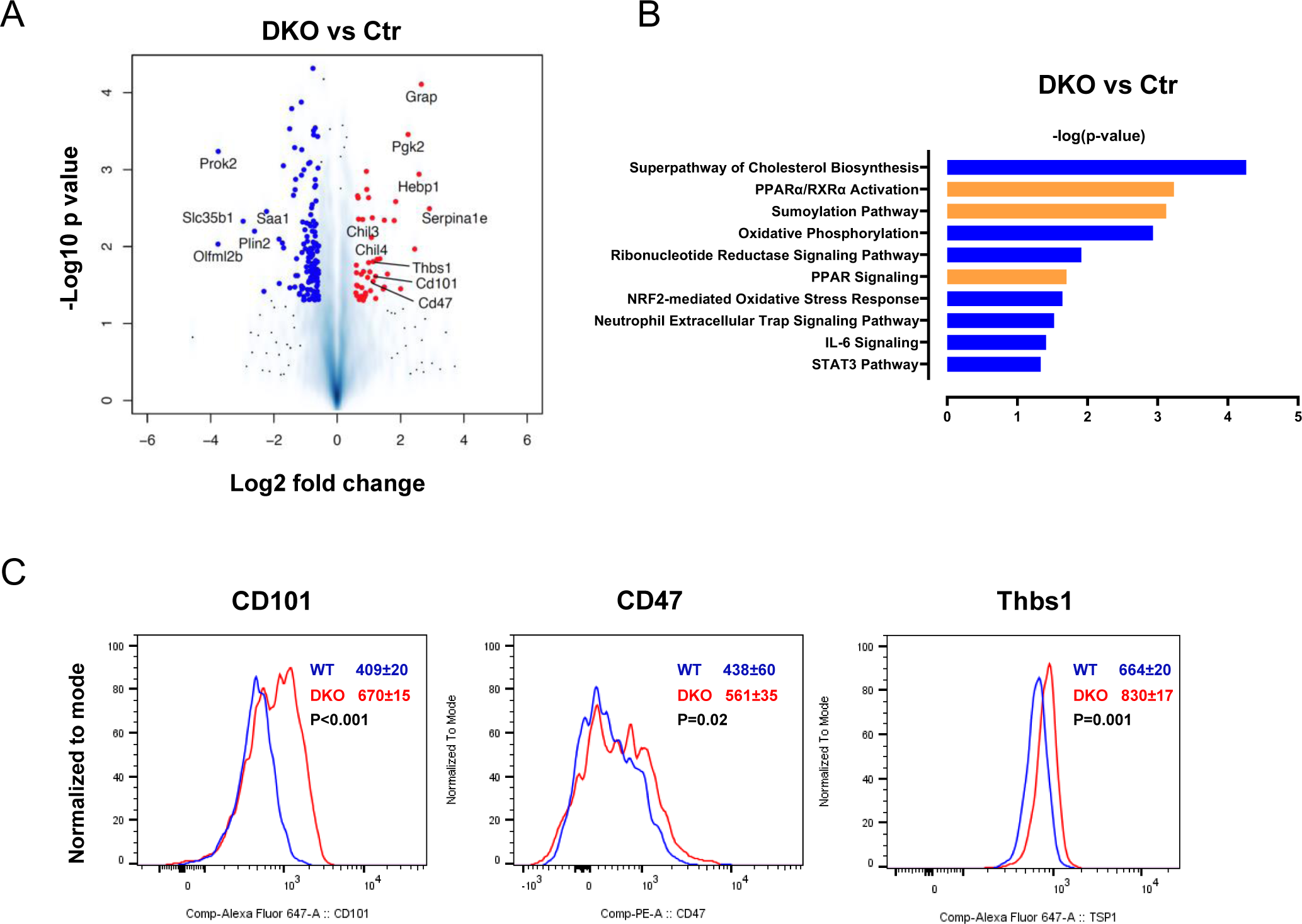
Upregulation of T cell suppressive proteins in circulating neutrophils from DKO-DSS mice. **(A-B)** Volcano plot of up- and down-regulated proteins (log2 fold change >0.7 or <-0.7, P value <0.001) in circulating neutrophil of DKO-DSS mice in comparison to those from WT-DSS mice and pathway analysis based on these altered proteins. Upregulated pathways in the DKO-DSS sample are in orange, whereas downregulated pathways are in blue. **(C)** Flow cytometry analysis of circulating neutrophils from DKO-DSS and WT-DSS mice for CD47, CD101 and Thbs1. Mean fluorescence intensity is shown as means±sem with P values (two-tailed Student’s *t*-test).

## Discussion

In this study, we investigated the roles of WNT5A and WNT5B in DSS-induced colitis and found that simultaneous inactivation of both *Wnt5a* and *Wnt5b* genes ameliorated DSS-induced colitis disease phenotypes. In addition, the DKO-DSS mice developed greater splenomegaly and stronger peripheral myeloid cell expansion than the control mice. Because neutrophil depletion and splenectomy abrogated the colitis-related phenotypic differences between the *Wnt5* DKO and control mice, together with the effect of *Wnt5* DKO on neutrophil development in the spleens, we conclude *Wnt5* DKO reprogramed neutrophils in the spleen to provide the protection for colitic injury.

Previous studies have suggested that neutrophils play an important role in IBD [35–37]. Neutrophil infiltration and its products (neutrophil extracellular traps, calprotectin, and proteases) correlate with the disease severity of IBD. It is not difficult to envision that neutrophil products, including reactive oxygen species, neutrophil extracellular traps, and proteases, can cause intestinal tissue damage. In addition, neutrophils produce a myriad number of cytokines that may recruit and activate neutrophils and modulate other immune cells. However, studies using experimental colitis models suggest the roles of neutrophils in intestinal inflammation and colitis are rather complex. Depletion of neutrophils led to the exacerbation of the colitis disease severity in not only DSS, but also dinitrobenzene sulphonic acid (DNBS) and T cell transfer colitis models [38, 39]. In our study, we also observed that neutrophil depletion in wild-type mice caused mild, but statistically significant, exacerbation of the colitis disease phenotypes. Neutrophils overlap with PMN-MDSCs in the function of immune cell suppression. MDSCs including PMN-MDSCs have been intensively investigated in the context of tumorigenesis, but also implicated in intestinal colitis inflammation regulation [40–42]. Given cytotoxic CD8^+^ T cells exacerbate colitis phenotypes, it is likely that the neutrophil functions as a double-edged sword; on one hand, it can cause tissue damage via its cytotoxic products and, on the other hand, it suppresses cytotoxic CD8^+^ T cells to dampen the damage. Our results that neutrophils from the DKO-DSS mice possess hyper-immune suppressive activity toward CD8^+^ T cells *in vivo* and *in vitro* and are largely responsible for the colitis-related and colonic CD8^+^ T cell suppression phenotypes caused by WNT5-deficiency support the importance of neutrophils with elevated immunosuppressive activity in curbing colitis disease development. In addition, the hypo-inflammatory characteristics of these DKO-DSS neutrophils revealed by scRNA-seq and proteomic analyses may also contribute to colitis protection.

Neutrophils are known to possess plasticity in their transcriptome, proteome, and function [43, 44]. The distinct transcriptomic and proteomic profiles of neutrophils produced in the absence of WNT5 from those in the presence of WNT5 or even PMN-MDSCs from tumor-bearing mice further demonstrate the high plasticity of this class of immune cells. The fact that *Wnt5* DKO leads to the development of hyper-immunosuppressive and hypo-inflammatory neutrophils to accentuate their beneficial roles in disease control (Fig. S7) suggests that neutrophil plasticity may be exploited for therapeutic benefits. Namely, neutrophils may be reprogrammed to possess characteristics that are less harmful to tissues and more beneficial for disease control.

It remains unclear how the lack of WNT5 reprograms neutrophils, which presumably occurs during splenic EMH, which is likely responsible for the splenomegaly phenotype. Chronic inflammation, including chronic intestinal inflammation, causes hematopoietic stress. This stress can trigger EMH, in which hematopoietic stem cells are mobilized to sites outside the bone marrow, the primary site for adult hematopoiesis, to expand hematopoiesis often in the spleen [43, 45–47]. Splenic EMH occurs with enlarged spleens, which are observed both in mice subjected to the experimental colitis model and in patients with Crohn’s disease [43, 48]. The splenic EMH often skews toward myelopoiesis, leading to excessive production of myeloid cells including neutrophils [49]. Thus, the peripheral myeloid cell expansion in *Wnt5 DKO* mice was likely the result of increased splenic EMH. This conclusion is supported by the observation that *Wnt5* DKO resulted in increased hematopoiesis in the spleens and reduced hematopoiesis in the bone marrow of DSS-treated mice. WNT5A has been implicated in bone marrow hematopoiesis and myelopoiesis, which are largely consistent with the bone marrow phenotypes of Wnt5 DKO[14, 50, 51]. The strong effect of *Wnt5* DKO on EMH is unexpected. We postulate that these WNT5 proteins may act directly on the hematopoietic stem/progenitor cells and/or myeloid cells or indirectly via altering the microenvironments of myelopoiesis. These important questions need to be addressed in future studies.

## Material and Methods

### Mice

*Rosa^creERT2^*, *Wnt5a^flox/flox^*, and *Wnt5b^-/-^* mice were purchased from Jackson Laboratory. *Wnt5a/5b* double knockout mice were created by crossing *Wnt5a^flox/flox^* mice with *Rosa^creERT2^* and *Wnt5b^-/-^* mice. All mice were age- (8 weeks old at the start of experiments) and sex-matched (both male and female mice were used). Littermate controls were used where appropriate. Mice were maintained in specific pathogen-free animal facilities at Yale University. The mice were housed under 12 hr light/dark cycles with free access to food and sterile water. All experiments were performed in accordance with guidelines from the Institutional Animal Care and Use Committee at Yale University.

### DSS-induced Colitis

To induce colitis in WT, *Wnt5b^+/-^*, *Wnt5b^-/-^*, *Rosa^creERT2^ Wnt5a^flox/flox^ Wnt5b^-/-^*, and *DKO* mice, 100mg/kg mouse body weight Tamoxifen (Sigma T-5648) dissolved in corn oil (Sigma C-8267) was delivered to mice at 8-week old through i.p injection, mice were then kept for DSS treatment until 13-week old to eliminate the side effect of Tamoxifen. 2% (36,000-50,000 M.Wt.) DSS (MP Biomedicals) was then delivered in drinking water for 7 days. Body weight loss as a clinical sign of colitis was recorded daily. The disease activity index including the presence of blood in stool, stool consistency, and rectal prolapse was calculated daily as has been previously described. Each mouse was scored from 0-4 according to the following criteria. Bleeding. 0: Normal (hemoccult negative, no visible blood in stool); 1: Hemoccult positive (hemoccult positive, no visible blood in stool); 2: Slightly visible blood in stool (hemoccult positive, visible blood in stool with reddish hue upon smear); 3: Visible blood in stool (hemoccult positive, obvious blood in stool, but no incrustation around anus); 4: Gross bleeding (fresh extensive blood around anus or encrusted on fur). Stool Consistency. 0: Normal (well-formed pellet, solid); 1: Soft (well-formed pellet, soft); 2: Pasty (semi-formed pellet, readily becomes paste upon handling); 3: Loose (poorly formed pellet, readily becomes paste upon handling); 4: Diarrhea (no pellet formation, and/or liquid stools). Rectal prolapse. 0: None; 1: Signs of prolapse; 2: Clear prolapse; 3: extensive prolapse. Final scores were determined by summing these three individual scores. Colonic lengths were measured immediately after colons were excised.

### Splenectomy Surgery

The surgery was performed three days before DSS treatment. The mice were under anesthesia and the abdominal cavity was opened. The spleen was carefully removed before splenic vessels were cauterized. For sham surgery, the abdomen was opened but the spleen was not removed.

### Colon Histology

H&E staining of colon tissues was performed at the Comparative Pathology Research core at Yale School of Medicine. Briefly, colons were excised, flushed with PBS, opened longitudinally, arranged in Swiss rolls, and fixed in 4% buffered formalin overnight. They were then embedded in paraffin, sectioned (5 μm), and stained with hematoxylin/eosin (H&E) according to standardized protocols. Slides were imaged using an Aperio eSlide Scanner (Leica Biosystems).

### Neutrophil Depletion *In Vivo*

Depletion of Ly6G^+^ neutrophils was performed by i.p. injection of the anti-Ly6G mAb (1A8, BioXCell BE0075) at a dose of 500 μg in 200 μL PBS per mouse. anti-Ly6G mAb was delivered every day starting from day -1 to day 6 of DSS treatment. Control mice received corresponding isotype mAb (BioXCell BE0089).

### Preparation of Single Cell Suspension From Tissues

#### Preparation of mouse colonic cells

Mouse colons were surgically excised, flushed with PBS, then opened longitudinally and cut into approximately 0.5-cm-long pieces. The colon tissue was washed again with 10 ml PBS followed by incubation in 20 ml HBSS buffer containing 2% FBS, 5mM EDTA and 1mM DTT in 50 ml canonical tube, at 37D°C with constant shaking at 180 rpm for 20 min to release epithelial cells. The colon tissues were then washed with 10 ml HBSS buffer before being digested in 10 ml 1640 buffer containing 2% FBS, collagenase type II (2mg/ml), collagenase D (0.5mg/ml), DispaseII (3mg/ml) and DNase I (0.1mg/ml) at 37D°C with constant shaking at 180 rpm for 30 min in a 15ml tube. After digestion, the digested cells and remaining tissue fragments were vortexed intensely for 20 s before being passed through a 40 μm cell strainer. The filtered cell suspension was centrifuged at 600g, 4D°C for 5 min, then resuspended in 1 ml Facs buffer (2% FBS, 2mM EDTA in DPBS).

#### Preparation of mouse splenocytes

Splenocytes were collected through mashing the spleen by using the plunger end of the syringe in 1 ml Facs buffer through placing the spleen into the cell strainer. Splenocytes were then filtered through a 40μm cell strainer. Cells were centrifuged for 5 min at 500*g*. The red blood cells were then lysed with 5 ml ACK lysis buffer for 3 min at room temperature. After centrifugation, cells were washed twice with Facs buffer and resuspended in Facs buffer.

#### Preparation of mouse bone marrow cells

BM cells were flushed from the femur and tibia bones with 15 ml Facs buffer and filtered through a 40_μ_m cell strainer. Cells were centrifuged for 5 min at 500*g*. Red blood cells were lysed with 1 ml ACK lysis buffer for 3 min at room temperature and washed twice with Facs buffer and resuspended in Facs buffer.

#### Preparation of mouse peripheral blood cells

Peripheral blood (600–800µl) was collected by retro-orbital bleeding in tubes with lithium heparin (BD 365965). Red blood cells were lysed by resuspension in 10 ml ammonium-chloride-potassium (ACK) lysis buffer (Sigma R7757) for 5 min at room temperature. Cells were washed twice with 10 ml Facs buffer before being resuspended in Facs buffer.

### Flow Cytometry

Single-cell suspensions were analyzed or sorted according to standard protocols using a BD FACS LSRII and BD FACS Aria III (BD Biosciences). Flow cytometry was performed as previously described [52]. Cells in single-cell suspension were stained with live-Dead dye (ThermoFisher L34957) for 20 min on ice in the dark. Cells were then fixed with 2% PFA (Santa Cruz, sc-281692). After being washed with a flow cytometry staining buffer (eBioscience, 00-4222-26), cells were first stained with Fc block (BD) for 10 minutes (except for GMP staining) followed by antibodies for cell-surface markers for 1 hour on ice in the dark. The cells were then washed, pelleted and resuspended in the flow cytometry staining buffer for flow cytometry analysis. The absolute number of cells was counted by using CountBright™ Absolute Counting Beads (Invitrogen C36950), according to the manufacturer’s instruction. Data were analyzed by flowjo.

### Sample preparation for Single-cell RNA-seq

Cells for single-cell RNA seq were sorted into PBS containing 0.04% BSA following the 10× Genomics protocol. Cell viability and counting were evaluated with trypan blue by microscopy, and samples with viabilities >90% were used for sequencing. Libraries were prepared using the Single Cell 5′ Library Kit V2 (10× Genomics). Transcriptome profiles of individual cells were determined by 10× Genomics-based droplet sequencing and libraries were sequenced with paired-end reads on an Illumina NovaSeq 6000 (Illumina).

### Single-cell RNA-seq analysis

The sequencing reads of the single-cell RNA-seq (scRNA-seq) was mapped to mouse reference (mm10) followed by quantification of transcript expression using cellranger V7.0.0. The single-cell RNA-seq data analysis was performed using Seurat v4.0.1 R package, including cell type stratification and comparative analyses between different experimental conditions (DKO, WT)[53]. In the quality control (QC) analysis, poor-quality cells with < 250 (likely cell fragments) or > 5,000 (potentially doublets) unique expressed genes were excluded. Cells were removed if their mitochondrial gene percentages were over 10% which indicates poor cell viability. The doublets from the scRNA-seq data were evaluated by DoubletFinder and were removed in the downstream analyses[54]. The data was first merged, normalized, and scaled with default settings in Seurat, followed by principal component analysis (PCA) for dimensionality reduction. We retained 30 leading principal components for further visualization and cell clustering. The Uniform Manifold Approximation and Projection (UMAP) algorithm was used to visualize cells on a two-dimensional space. Subsequently, the share nearest neighbor (SNN) graph was constructed by calculating the Jaccard index between each cell and its 20-nearest neighbors, which was then used for cell clustering based on Louvain algorithm (with a resolution of 0.3). Each cluster was screened for marker genes by differential expression analysis based on the non-parametric Wilcoxon rank sum test. Based on checking the expression profile of those cluster-specific markers, we identified 8 distinct cell types (including T cells, neutrophils, Mono/Macro, myeloid progenitor cells, two types of B cells, and two types of DCs). In the downstream analysis, we focused on the neutrophil populations which consists of four subclusters. Similarly, we examined the marker profiles of each subcluster to distinguish the functional roles of different neutrophil subtypes. We then specifically identified differentially expressed genes (DEGs) among each sample. Top representative DEGs were visualized using heatmaps or volcano plots.

### Public data collection and integration

Two independent tumor spleen neutrophil scRNA-seq datasets (GSE139125 and GSE163834) and tumor infiltration neutrophil scRNA-seq dataset (GSE213861) were downloaded from Gene Expression Omnibus (GEO) database. Raw sequencing reads were processed and analyzed using cellranger as mentioned above. Seurat was utilized for the down-stream analysis. Batch effect caused by the datasets were assessed and removed using harmony package in R[55]. DEGs among the clusters were identified using FindAllMarker function in Seurat package, where the dataset or batch was regarded as a covariate and was then adjusted. Over-representative pathways of the DEG in the clusters were evaluated by ShinyGO (http://bioinformatics.sdstate.edu/go/).

### Mass spectrometry sample preparation and data procession

The cell lysis buffer 10 M urea containing the cOmplete™ protease inhibitor cocktail (Roche, #11697498001) was used for protein extraction. A VialTweeter device (Hielscher-Ultrasound Technology)[56, 57] was used for sonication at 4°C for two cycle (1 min per cycle), and then centrifuged at 20,000 x g for 1 h to remove the insoluble materials. To break the disulfide bridge and “cap” the cystine, the reduction and alkylation were conducted with 10 mM Dithiothreitol (DTT) for 1 h at 56 °C and then 20 mM iodoacetamide (IAA) in dark for 45 min at room temperature. The samples were diluted by 100 mM NH4HCO3 and digested with trypsin (Promega) at ratio of 1:20 (w/w) overnight at 37 °C. The digested peptides purification was performed on C18 column (MarocoSpin Columns, NEST Group INC) and 1 µg of the peptide was injected for mass spectrometry analysis.

The samples were measured by data-independent acquisition (DIA) mass spectrometry method as described previously[58–60]. The Orbitrap Fusion Tribrid Lumos mass spectrometer (Thermo Scientific) instrument coupled to a EASY-nLC 1200 systems (Thermo Scientific, San Jose, CA) was used. The data acquisition was performed with 150-min gradient and 300 nL/min flow rate with the temperature controlled at 60 °C using a column oven (PRSO-V1, Sonation GmbH, Biberach, Germany). All the DIA-MS methods consisted of one MS1 scan and 33 MS2 scans of variable isolated windows with 1 m/z overlapping between windows. The MS1 resolution is 120,000 at m/z 200. The MS1 full scan AGC target value was set to be 2E6 and the maximum injection time was 50 ms. The MS2 resolution was set to 30,000 at m/z 200 and the normalized HCD collision energy was 28%. The MS2 AGC was set to be 1.5E6 and the maximum injection time was 50 ms. The default peptide charge state was set to 2. Both MS1 and MS2 spectra were recorded in profile mode. DIA-MS data analysis was performed using Spectronaut v16[61–63] with directDIA algorithm by searching against the SwissProt downloaded mouse fasta file. The oxidation at methionine was set as variable modification, whereas carbamidomethylation at cysteine was set as fixed modification. Both peptide and protein FDR cutoffs (Qvalue) were controlled below 1% and the resulting quantitative data matrix were exported from Spectronaut. All the other settings in Spectronaut were kept as Default.

### *In vitro* T cell suppression assay

CD8^+^ T cell preparation: Splenocytes were freshly isolated from mice. Red blood cells were lysed using RBC lysis buffer before being magnetically sorted using the CD8^+^ T cell isolation kit (Catalog # 130-096-543, Miltenyi Biotec). Neutrophil preparation: Splenocytes were freshly isolated from mice. Red blood cells were lysed using RBC lysis buffer before being magnetically sorted using the Anti-Ly-6G MicroBeads UltraPure (Catalog # 130-120-337, Miltenyi Biotec). T cells were distributed into the round bottom 96 well plates at 3000 cells per well and stimulated by CD3/CD28 mAb-coated beads (ThermoFisher Scientific) at a ratio of 10:1. Neutrophils were seeded with T cells at a concentration of 2:1. 48 hours after co-culture, T-cell activation was measured by flow cytometry.

### Statistical analysis

Comparisons of means between two groups and multiple groups were tested by unpaired, two-tailed *t* test and Two-way ANOVA-test using Prism 9.2.0 software (GraphPad). Statistical tests used biological replicates. *p < 0.05, **p < 0.01 and ***p < 0.001 were all considered statistically significant.

---

**Figure S1.**
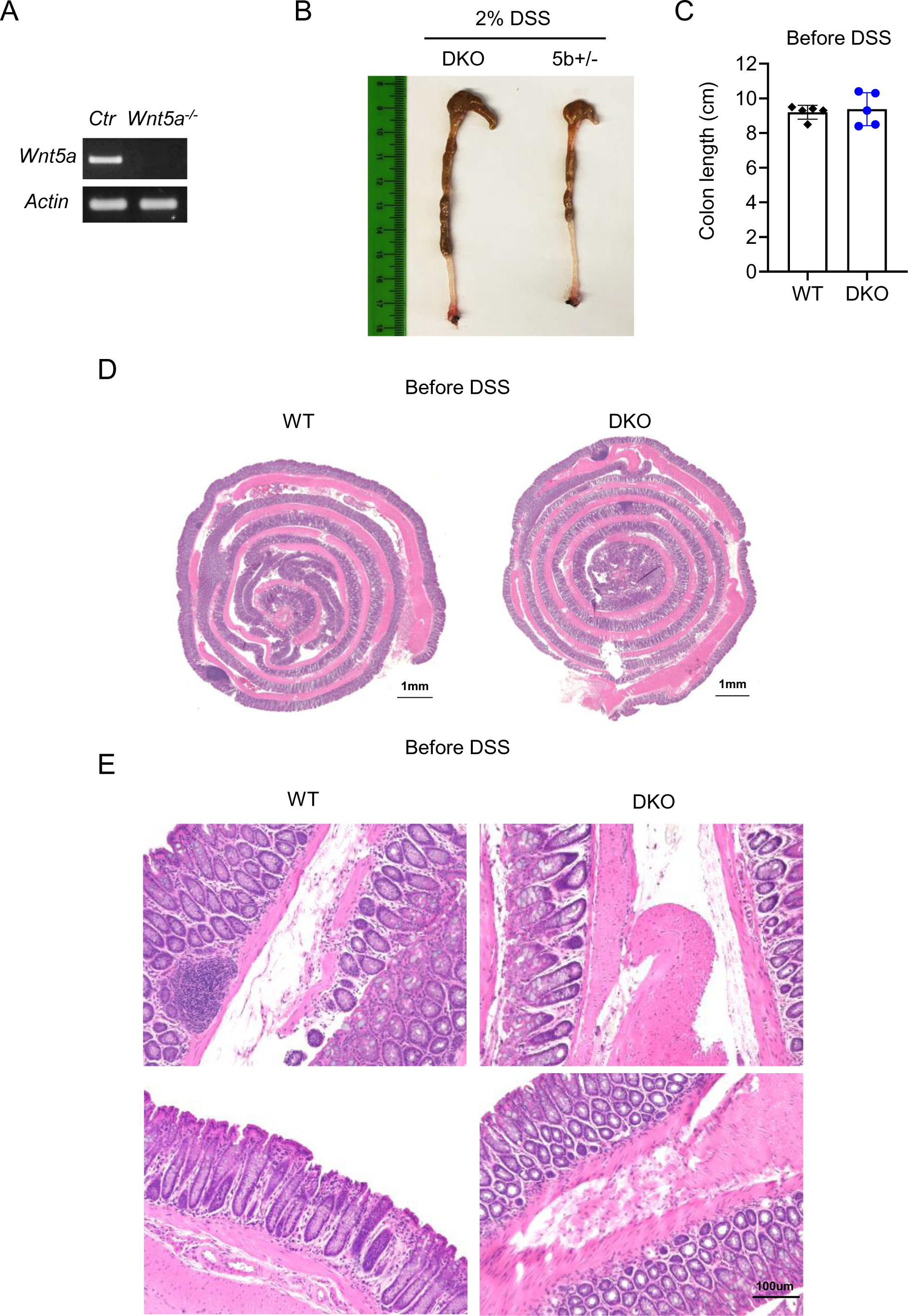
Loss of WNT5 protects mice from DSS-induced colitis. **(A)** RT-PCR validation of *Wnt5a* knockout efficiency. **(B)** Representative colons from DKO and control mice after 7-day DSS treatment are shown. **(C-E)** Colon length and H&E sections of colons from the DKO and control mice collected before DSS treatment are shown. Data in **(C)** are presented as means±sem. Each datum point represents one mouse.

**Figure S2.**
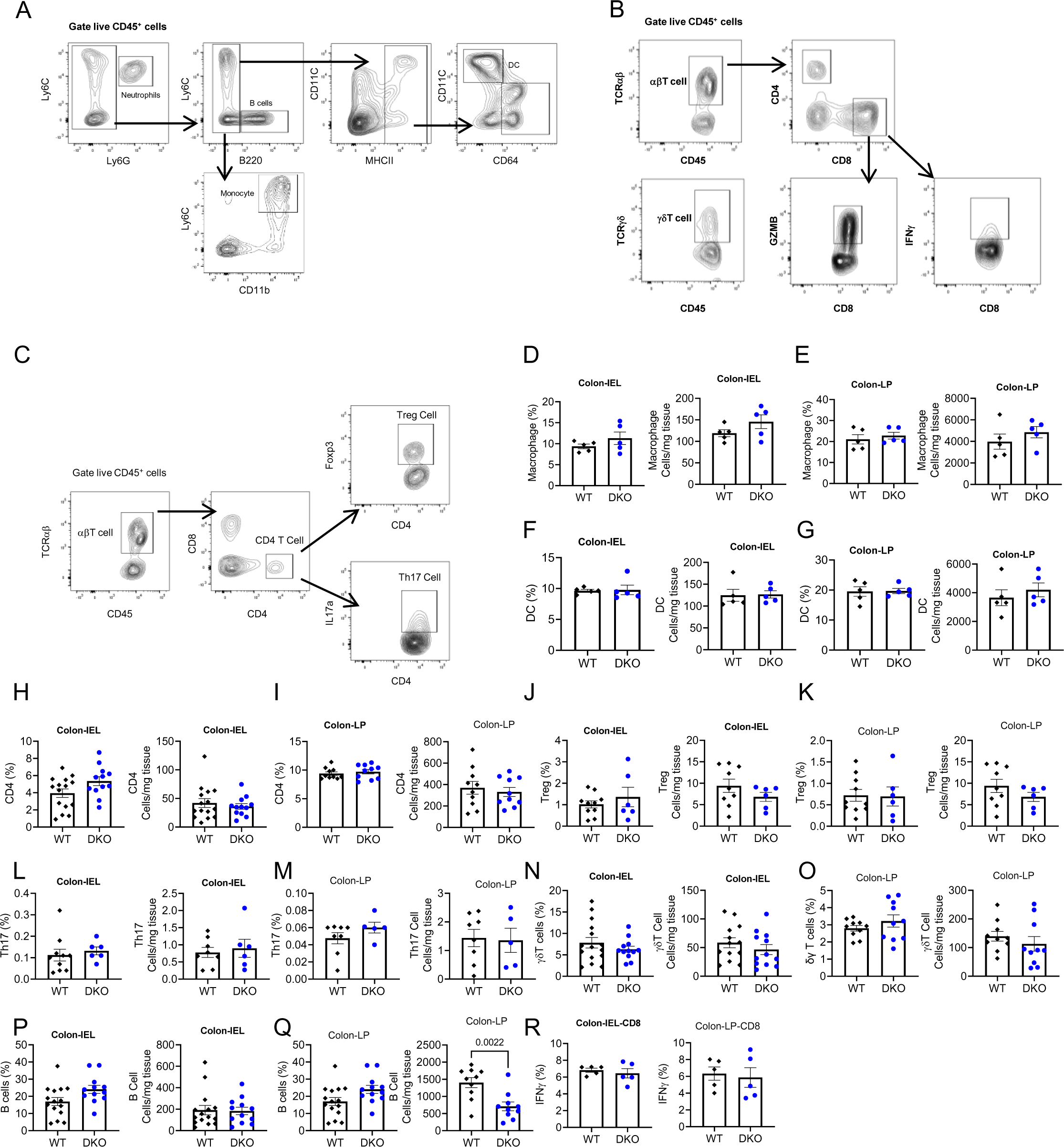
Immune cell compositions in the colons of *Wnt5* DKO and control mice. **(A-C)** Flow cytometry gating strategies for each population are shown. (**D-R**) Mice were treated as in Fig. 1, and percentage (in CD45^+^) and absolute number of various immune cell populations were determined by flow cytometry. Data are shown as means±SEM with P value (two-tailed Student’s *t*-test). Each datum point represents one mouse.

**Figure S3.**
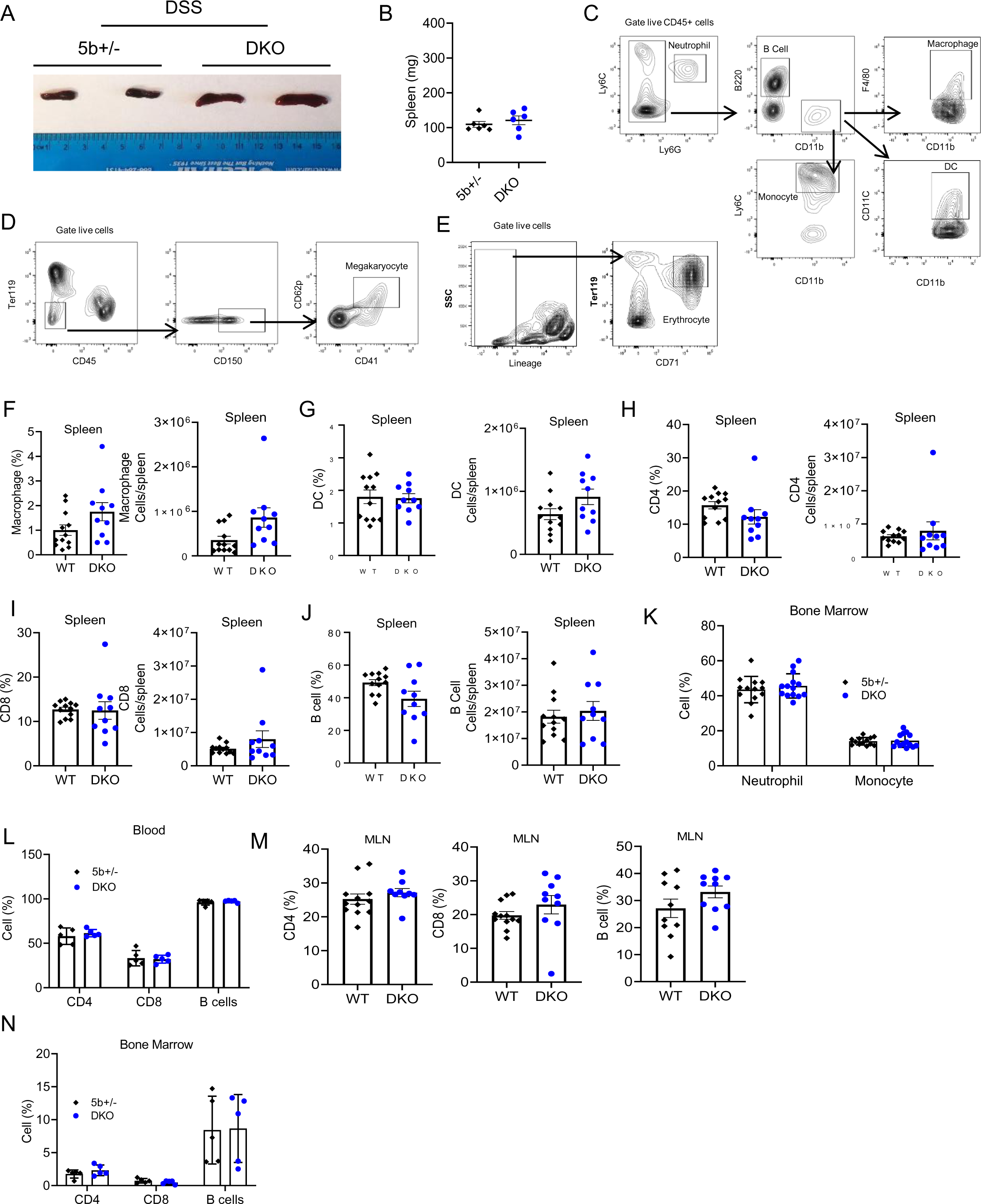
Colitic *Wnt5* DKO mice develop splenomegaly. **(A)** Representative images of spleens from the DKO and control mice at Day 7 of DSS treatment. **(B)** Spleen weight of *Wnt5* DKO and control mice without DSS treatment. **(C-E)** Flow cytometry gating strategies for the CD45^+^-gated cells from spleens. The same gating strategy was used for flow cytometry analysis of blood and MLN cells. (**F-N**) Mice were treated as in Figure 1 and the percentage and the absolute number of immune cells were determined by flow cytometry. Results in **(B, F-N)** are shown as means±sem with P value (two-tailed Student’s *t*-test). Each datum point represents one mouse.

**Figure S4.**
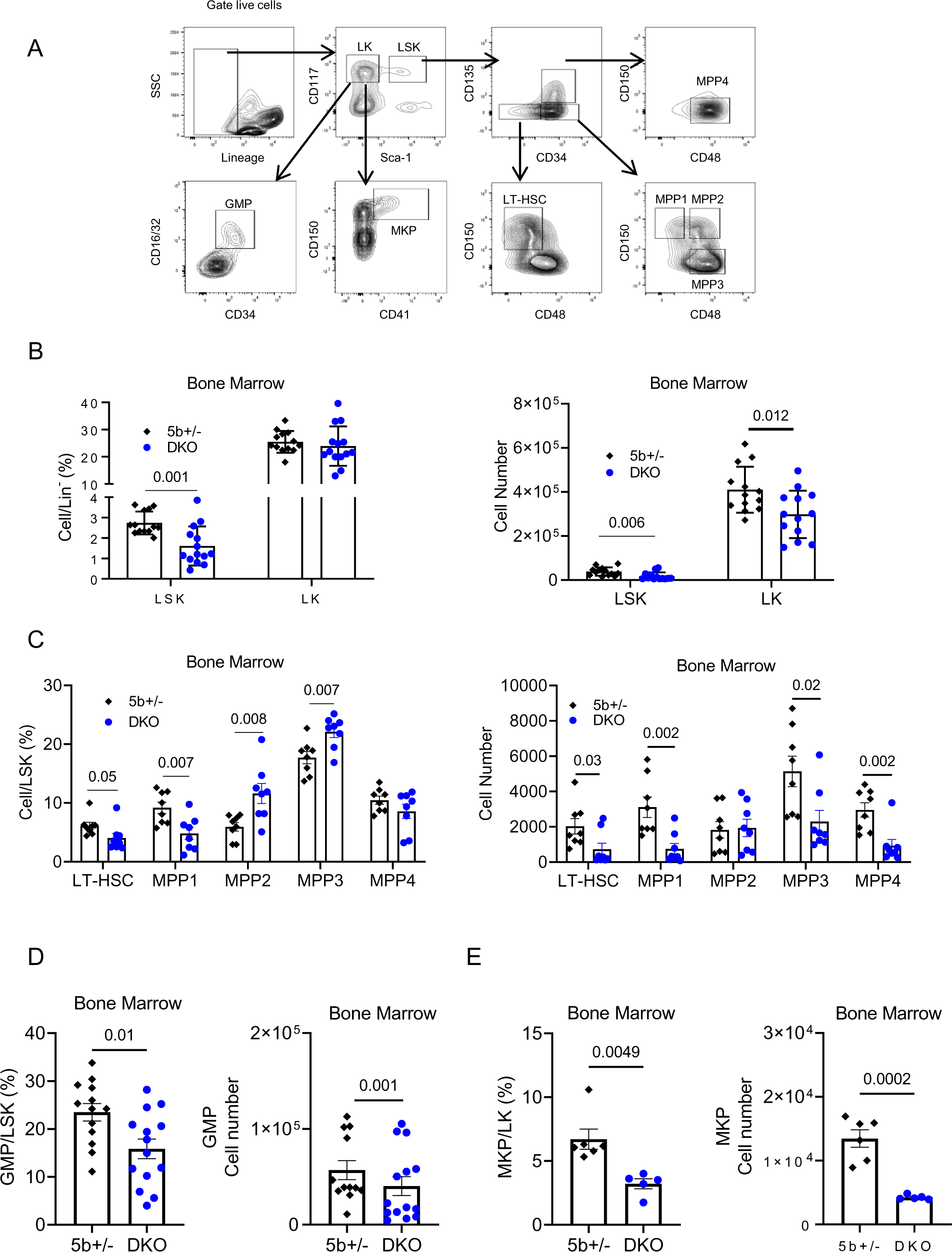
*Wnt5* DKO decreases bone marrow hematopoiesis upon colitis induction. **(A)** Gating strategies for flow cytometry analysis of hematopoietic cell stem/progenitor populations during hematopoiesis in the bone marrow and spleen are shown. **(B-E)** Mice were treated as in Figure 1 and the percentage (in Lineage-) and the absolute number of LK and LSK **(B)**, percentage (in LSK) and absolute number of LT-HSC, MPP1-4 **(C)**, percentage (in LK) and absolute number of GMP **(D)** and MKP **(E)** in the bone marrow of the mice described in Fig. 4 were determined by flow cytometry. Results in **(B-E)** are shown as means±sem with P values (two-tailed Student’s *t*-test). Each datum point represents one mouse.

**Figure S5.**
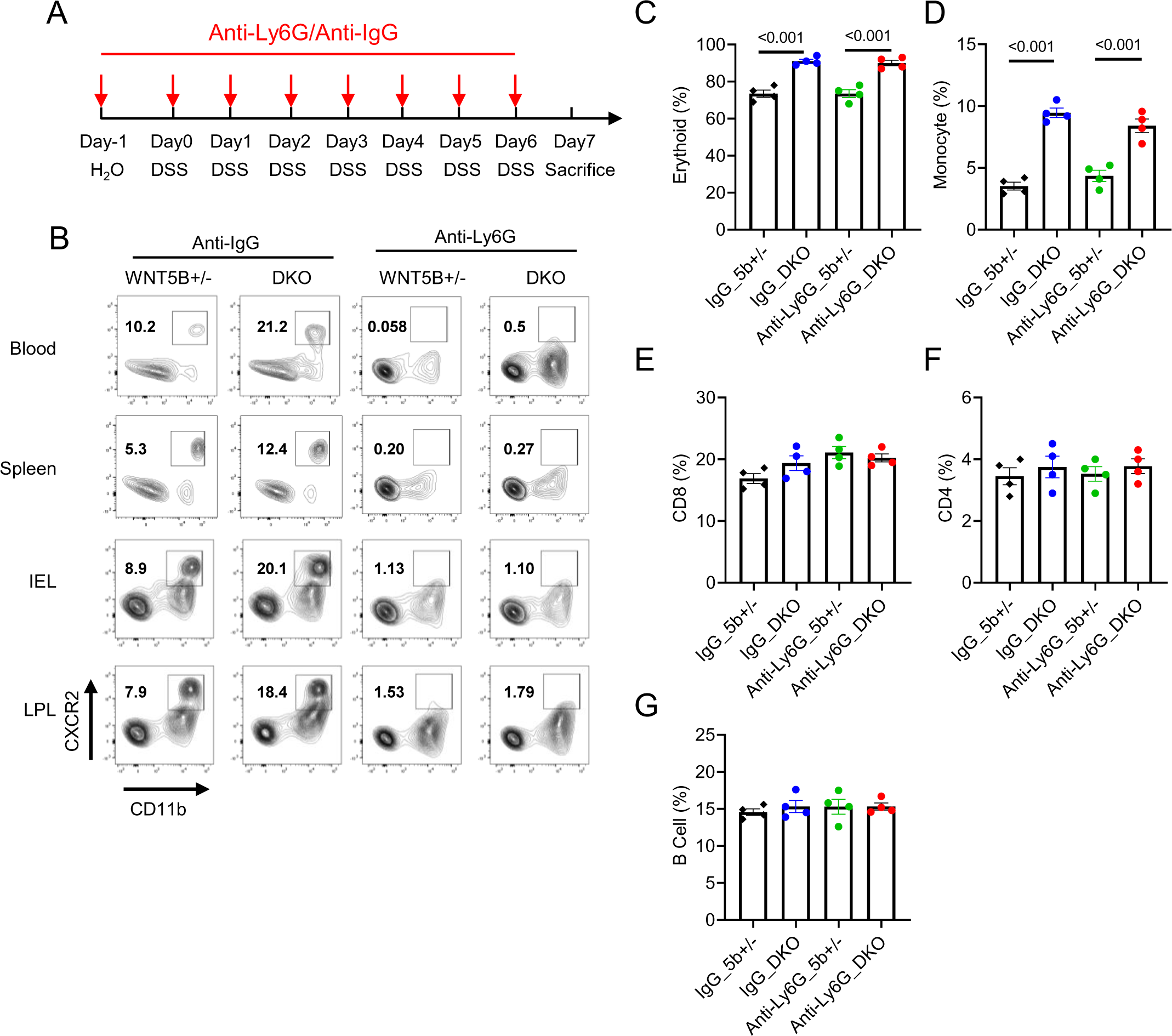
Neutrophil depletion procedure and efficiency. **(A)** Schematic representation of the procedure used to deplete neutrophils in the *Wnt5* DKO and control mice subjected to DSS-induced colitis. (**B-G**) *Wnt5* DKO and the control mice were treated with DSS and subjected to neutrophil depletion as in **A**. Neutrophil depletion efficiency in various tissues is shown in **(B)**. The frequencies of erythrocytes and monocytes in the spleen are shown in **(C)** and **(D)**. The frequencies of CD8^+^ T cells, CD4^+^ T cells, and B cells in the colon are shown in **(E-G)**. Data are presented as means±SEM with P values (two-tailed two-way ANOVA). Each datum point represents one mouse.

**Figure S6.**
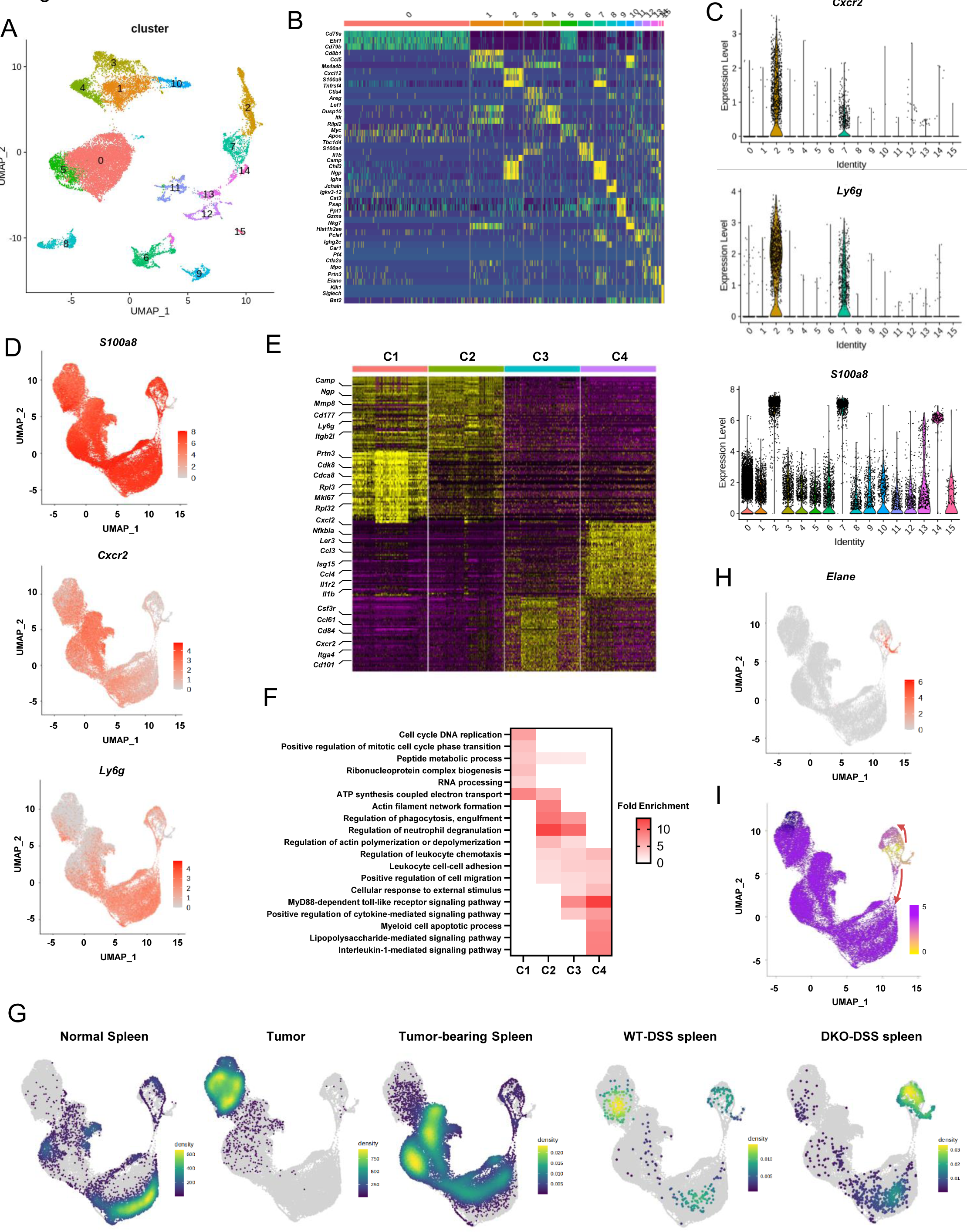
Single-cell RNA sequencing of splenic cells. **(A)** UMAP plot of all of the splenic cells (21,745 in total) from WT-DSS and KO-DSS mice that were analyzed by scRNA-seq and passed quality control. **(B)** The expression of cell type-marker genes in each cluster identified in **(A)**. **(C)** The expression of neutrophil-marker genes in each cluster identified in **(A)**. **(D)** The expression of neutrophil-marker genes in cells was shown in Fig. 6E. **(E)** Gene expression heat map of the neutrophils shown in Fig. 6E. Selected genes are marked, and the full list of the genes was described in Table S2. **(F)** Gene Ontology analysis of DEGs for each subcluster in Fig. 6E. **(G)** Point density-plot visualization of neutrophil distributions under different conditions. The color represents the density of overlapped neutrophils in local regions. **(H)** The expression of *Elane* in the neutrophils was shown in Fig. 6E**. (I)** Pseudotime plot as in Fig. 6F, but in a compressed scale, to illustrate two developmental trajectories from the *Elane*^+^ cells.

**Figure S7.**
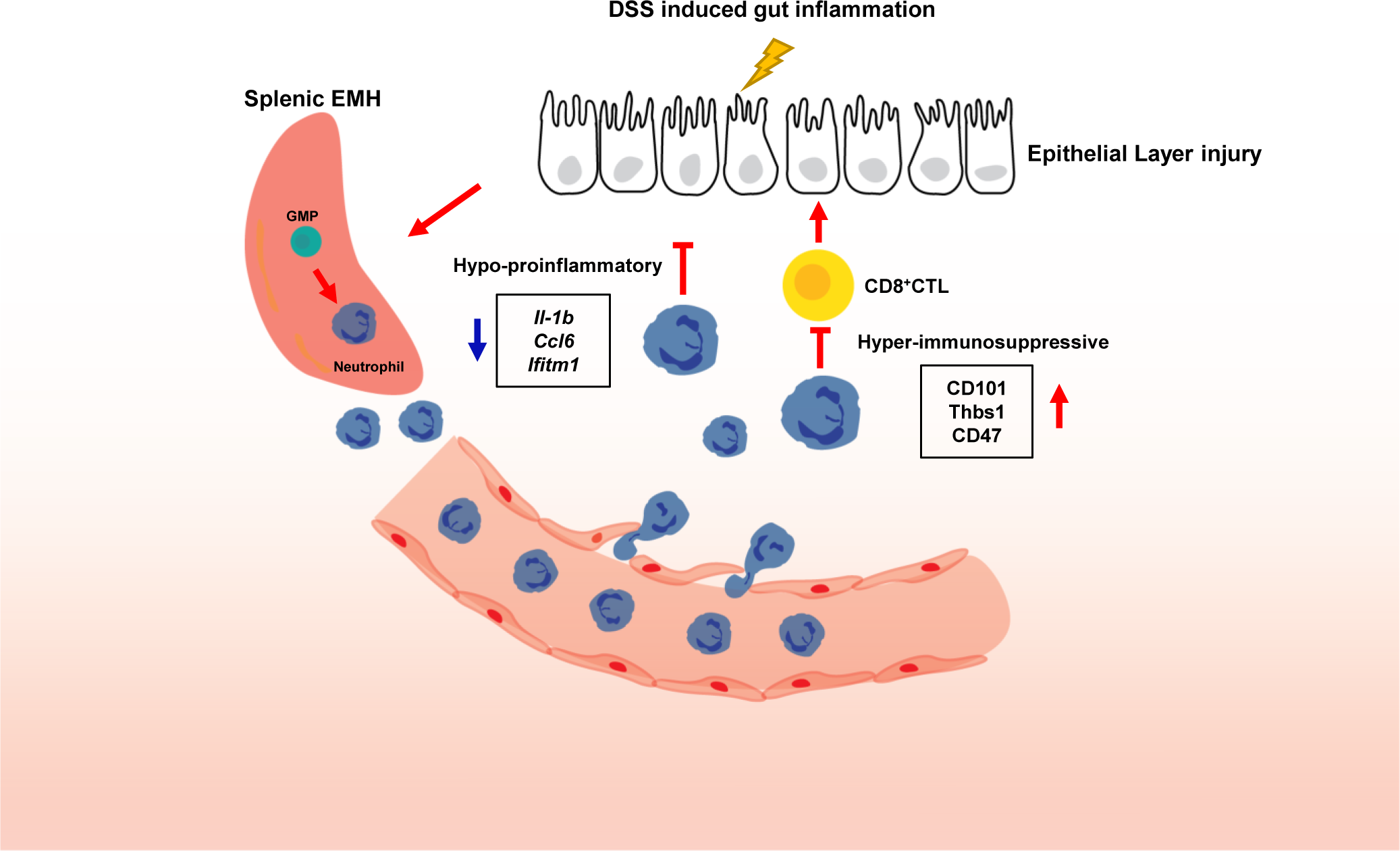
Graphic summary of the role of neutrophils produced in WNT5 deficient mice in the protection of DSS-induced colitis. The loss of WNT5 proteins in DSS-treated mice leads to splenomegaly, extramedullary hematopoiesis, and generation of neutrophils in spleens that show strong suppression of CD8^+^ cytotoxic T cells and reduced expression of pro-inflammatory cytokines. These neutrophils protect mice from DSS-induced colitis.

## Notes

### Competing Interest Statement

The authors have declared no competing interest.

